# Early-life stress facilitates the development of Alzheimer’s disease pathology via angiopathy

**DOI:** 10.1101/2020.04.27.062729

**Authors:** Tomoko Tanaka, Shinobu Hirai, Masato Hosokawa, Takashi Saito, Hiroshi Sakuma, Takaomi Saido, Masato Hasegawa, Haruo Okado

**Affiliations:** Laboratory of Neural Development, Department of Psychiatry & Behavioral Science, Tokyo Metropolitan Institute of Medical Science, Tokyo, Japan; Dementia Research Project, Department of Brain & Neuroscience, Tokyo Metropolitan Institute of Medical Science, Tokyo, Japan; Laboratory for Proteolytic Neuroscience, RIKEN Center for Brain Science, Saitama, Japan; Department of Neurocognitive Science, Institute of Brain Science, Nagoya City University Graduate School of Medical Science, Aichi, Japan; Child brain Project, Department of Brain & Neuroscience, Tokyo Metropolitan Institute of Medical Science, Tokyo, Japan

**Keywords:** Alzheimer’s disease, angiopathy, immune system, early life stress, amyloid-beta, cognitive function

## Abstract

**Background:** Alzheimer’s disease (AD), a progressive neurodegenerative disorder, is a serious social problem. Recently, several early-life factors have been associated with an increased risk of a clinical diagnosis of AD.

**Methods:** We investigated the involvement of early-life stress in AD pathogenesis using heterozygous the amyloid precursor protein (APP) mutant mice (*App^NL-G-F/wt^*) and wild-type (*App^wt/wt^*) mice. Maternal separation was used as an animal paradigm for early-life stress. Object location and fear conditioning tests were performed to measure cognitive functions, in addition to biochemical tests. Immunohistochemical analyses were performed after the behavioral tests.

**Results:** We found that maternal-separated *App^wt/wt^* mice showed narrowing of vessels and decreased pericyte coverage of capillaries in prefrontal cortex, while maternal-separated *App^NL-G-F/wt^* mice additionally showed impairment of cognitive function, and earlier formation of Aβ plaques and disruption of the blood–brain barrier. Severe activation of microglia was detected in the maternal-separated *App^NL-G-F/wt^* mice and maternal-separated *App^wt/wt^* mice. At the early stage, morphological changes and inflammatory responses were observed in the microglia of the maternal-separated *App^NL-G-F/wt^* mice and maternal-separated *App^wt/wt^* mice, as well as morphological changes in the microglia of the non-maternal-separated *App^NL-G-F/wt^* mice.

**Conclusions:** Microglia activation induced by maternal separation in combination with the APP mutation may impairs the vascular system, leading to AD progression. These findings therefore suggest that maternal separation causes early induction of AD pathology via angiopathy.

## Introduction

The development of abnormal physiology in association with increased age is observed in the progression of many disorders. Because daily habits and environmental stresses such as alcohol intake and diet have the major influence on life-span, rather than genetic factors (Christensen et al., 2006), acceleration of the aging process by such environmental factors is thought to increase both the risk of developing a disease and mortality from a disease. Alzheimer’s disease (AD) is one such disease that is thought to be influenced by environmental factors; it is characterized by progressive impairment of cognitive functions such as memory. It is considered that the acquired predisposition to AD increases with age, and that trauma during adulthood increases the likelihood of developing AD. Moreover, recent epidemiological research has suggested that stressful environmental factors during the developmental period, including mistreatment, neglect, and the loss of a parent, can also increase the likelihood of developing AD (Norton et al., 2011; Radford et al., 2017; Seifan et al., 2015). However, as the research behind these findings is generally the result of retrospective cohort studies, it is unclear whether early-life stress affects the onset and progression of AD.

Recently, vascular impairment, a postmortem pathological finding in the brain, was reported to occur before the onset of AD. Therefore, vascular impairment is considered as an initial symptom of AD (Iadecola, 2017; Montagne et al., 2016; Wen et al., 2004). The neurovascular unit (NVU) consists of neurons, endothelial cells, pericytes, microglia, and astrocytes. As the NVU is important for the extrusion of amyloid beta, an impairment in extraction can cause sporadic AD and cerebral amyloid angiopathy (Mawuenyega et al., 2010). In particular, the capillary vessels in the brain play an important role in the extrusion of amyloid beta. Dysfunction of the blood–brain barrier (BBB) and a decrease in amyloid beta clearance through the cerebral vascular system is observed in AD (Zlokovic, 2011). Degeneration of cells making up the BBB, such as pericytes, causes dysfunction of the BBB (Sweeney et al., 2016). However, it remains unclear whether early-life stress results in dysfunction of the vascular system.

In this study, we investigated whether maternal separation, which can be used as an early-life stress in rodents, facilitates the development of AD, with the purpose of understanding the causal association between the development of AD and early-life stress. After finding indications that maternal separation affects the development of AD, we next investigated the changes induced by maternal separation, paying particular attention to capillary vessel morphology in heterozygous the amyloid precursor protein (APP) mutant mice (*App^NL-G-F/wt^*) and wild-type (*App^wt/wt^*) mice. As a result, we established that maternal separation facilitates the development of AD. Furthermore, we found that impairment of capillaries, probably caused by abnormalities in microglia, leads to dysfunction of the BBB, thereby facilitating the development of clinical AD.

## Materials and methods

All experimental procedures were approved by the Animal Experimentation Ethics Committee of the Tokyo Metropolitan Institute of Medical Science (49040).

### Animals

The original lines of App mutant mice (*App^NL-G-F/NL-G-F^*) were established as a C57BL/6J congenic line (a genetic background strain) by repeated back-crosses, as described previously, and were obtained from RIKEN Center for Brain Science (Wako, Japan)(Saito et al., 2014). All experiments were performed with *App^NL-G-F/wt^* and *App^wt/wt^* that were born from *App^NL-G-F/wt^* mice x *App^NL-G-F/wt^* mice or *App^NL-G-F/wt^* mice x *App^wt/wt^* mice. *App^NL-G-F/wt^* mice with slow onset of pathology were used because it was important to investigate the process of decline in function. All mice were maintained under a 12:12 h light/dark cycle (lights on at 8:00 AM). All efforts were made to minimize the number of animals used and their suffering. All experiments, except for maternal separation, were performed in adolescence (postnatal day 45 (P 45)) or adulthood (P90–P120).

### Maternal separation

Maternal separation is usually used as a form of early-life stress in mice. Pups were separated from their mother for 3 hours (AM 9:00-12:00) daily from P2 to P15. The separated pups were taken to a different room and placed on a heat pat to maintain their body temperature. The control mice had no experience of maternal separation. The body weights were not significantly different between the non-maternal-separated and maternal-separated mice at several stages of development.

### Behavioral assessment

Male mice were allowed to habituate in the behavioral room for at least 1 hour before commencement of the behavioral test. Mice showing outlier behavior > 2 standard deviations from the mean or < 2 standard deviations from the mean were not analyzed by the behavioral test.

The object location test consisted of three phases. All phases were performed under a light intensity of 10 lux. On day 1, the animals were placed in an empty square box (50 cm × 50 cm) for 10 minutes. On day 2, the animals were placed in the same box for 10 minutes with two same identical unfamiliar objects. On day 3, the animals were placed in the same box, with one of the two items being displaced to a novel location in the area. The discrimination index was calculated as the ratio of the time spent exploring the novel place to that spent exploring the familiar place: discrimination index = (novel place exploration time – familiar place exploration time)/(novel place exploration time + familiar place exploration time). On each day, the exploration time in each place was calculated automatically using a DVTrack video tracking system (Muromachi, Tokyo, Japan).

The fear conditioning test consisted of three phases. On day 1, the animals were placed in a triangular box for 5 minutes under a light intensity of 30 lux. On day 2, the animals were placed into a square box with a stainless-steel grid floor and were allowed to explore the box freely for 2 minutes. Subsequently, a tone, which served as the conditioned stimulus (CS), was presented for 30 sec. During the last 1 sec of CS presentation, a 0.75 mA electric shock was applied, which served as the un-conditioning stimulus (US). Two more CS-US parings were presented with a 1 minutes inter-stimulus interval under a light intensity of 100 lux. On day 3, the animals were placed into the same square box as used on day2 for 5 minutes under a light intensity of 100 lux. On day 4, the animals were placed in the same triangular box as used on day 1 for 10 minutes and were allowed to explore the box freely for 2 minutes. Subsequently, CS was presented for 30 sec. Seven more CS were presented with a 1 minutes inter-stimulus interval under a light intensity of 100 lux. In each test, the percentage of freezing time and distance travelled (cm) were calculated automatically using ImageFZ software (O’Hara & Co., Tokyo, Japan). Freezing was taken to indicate that the mice remembered the box (context) or tone (cue) in which they were exposed to the electrical shock.

### Immunohistochemistry

At P45 or P120, male and female mice were anaesthetized with isoflurane and then sequentially perfused through the heart with PBS followed by 4% paraformaldehyde (PFA) in PBS. Their brains were post-fixed, cryoprotected in 20% sucrose at 4°C, and embedded in OCT compound and frozen. Coronal sections of mouse brain were then cut using a cryostat (20 μm). Free-floating sections containing PFC were activated by HistVT one (Nakalai., Kyoto, Japan) at 70°C for 20 minutes, incubated in 0.4% Block-ase (DS Pharma Biomedical., Osaka, Japan) in PBS at room temperature for 20 minutes, and then incubated with primary antibody (Supplemental figure 1). All primary antibodies were diluted 1:500 in PBS containing 0.3% Triton X-100. The sections were then incubated with secondary antibody (Jackson., Maine, United State) (diluted 1:500 in PBS containing 0.3% Triton X-100) for 2 hours at room temperature. All sections types were incubated with DAPI (Nacalai Tesque., Kyoto, Japan) for nuclear staining. Fluorescence micrographs of the sections subjected to immunostaining procedures were captured and digitized with a FluoView® FV3000 confocal laser scanning microscope (Olympus., Tokyo, Japan). With the aim of quantifying the morphological change of microglia, skeleton analysis and particle analysis were performed by ImageJ(Fernández-Arjona et al., 2017; Xu et al., 2018).

### Luminex analysis assay

Following the sacrifice of male and female mice, mouse PFC fractions were dissociated with a tissue homogenizer in 100 μl of 0.5 × cell lysis buffer 2 diluted in PBS (R&D Systems., Minnesota, United State). Tissues were lysed at room temperature for 30 minutes, and debris was then removed by centrifugation. Magnetic luminex assays were performed using a Mouse Premixed Multi-Analysis Kit (R&D Systems., Minnesota, United State). Assay procedures were carried out following manufacturer’s instructions. Microplates were run on a Bio-Plex 200 system (Bio-Rad., California, United State).

### Corticosterone assay

Male and female mice were subjected to a novel cage stress (Rattray et al., 2009), in which each mouse was placed in isolation in a new cage with a smooth floor for 30 minutes at P45. Blood samples were taken before stress and immediately after stress, and centrifuged at 3,300 rpm to separate serum. Corticosterone concentrations were measured using an enzyme-linked immunosorbent assay (ELISA) (AssayPro., Missouri, United State).

### Statistical data analysis

Group data are presented as the mean ± SEM. The statistical significance of between-group differences was assessed by the Tukey–Kramer test, Dunnett’s test, or the paired t-test, using JMP software (SUS Institute., North Carolina, United State). The interaction between area and distance was assessed using the polynomial regression model. The statistical significance of differences between groups and over time was assessed by two-way repeated ANOVA followed by a *post-hoc* Holm’s test, using R software.

## Results

### Experimental design

Figure 1A shows the experimental schedule. To investigate whether stress in the early developmental stage is involved in the onset and/or progression of AD, we performed maternal separation for 3 hours daily during the period from P2 to P15 in *App^wt/wt^* and *App^NL-G-F/wt^* mice. The *App^NL-G-F/wt^* mice show an AD-like phenotype in the late-aged stage (Saito et al., 2014). We then performed behavioral tests and tissue sampling around P45 (adolescence) and P120 (adult).

**Figure 1.**
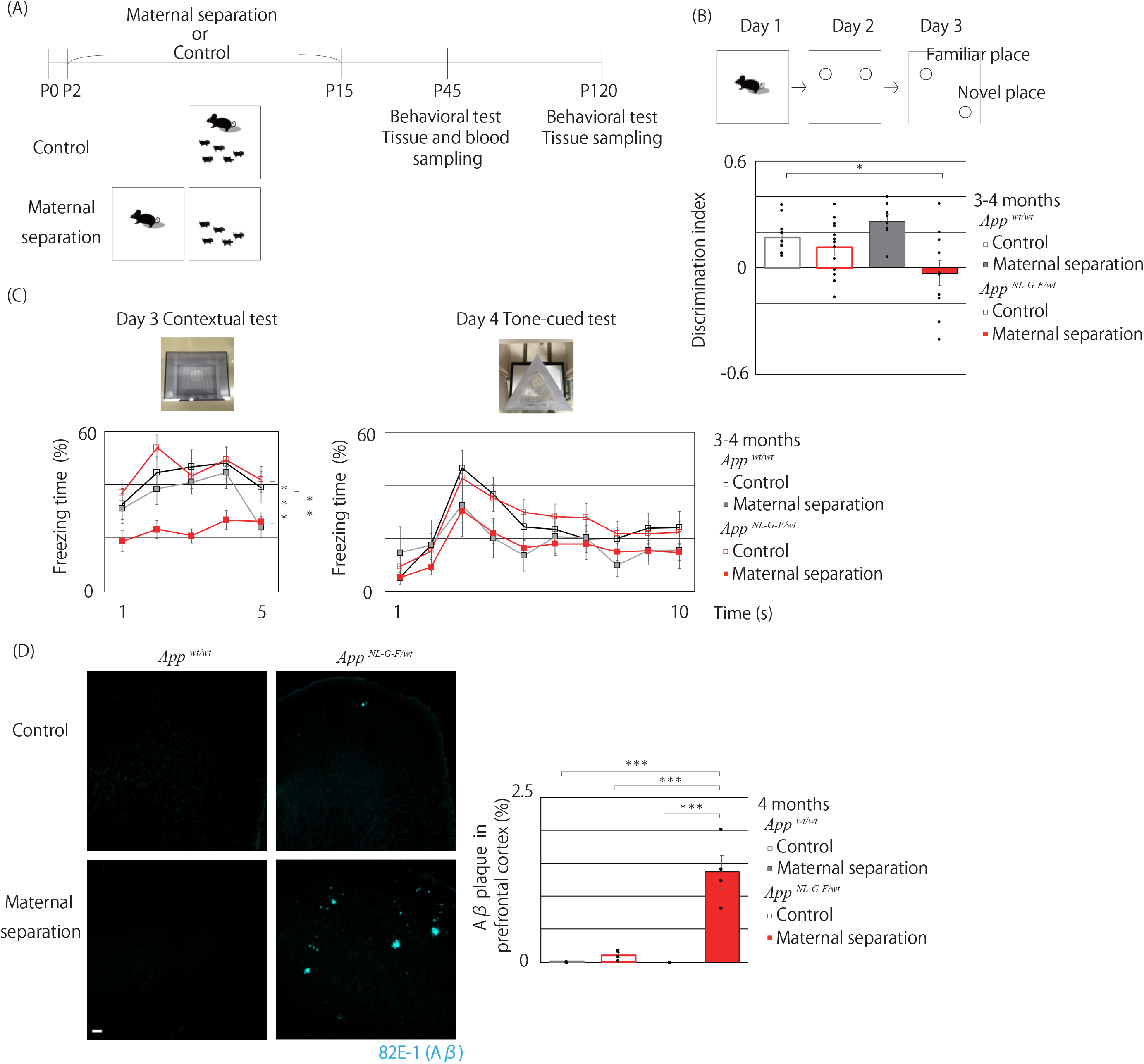
Maternal separation induces Alzheimer’s disease-like behavior and amyloid beta plaque formation. (A) Time schedule for maternal separation and other investigations. (B) Spatial memory was measured using an object location test. Behavioral layout of the object recognition test for the four experimental groups. Maternal-separated *App^NL-G-F/wt^* mice showed impairment of spatial memory. *p < 0.05 Maternal-separated *App^NL-G-F/wt^* mice vs non-maternal-separated *App^wt/wt^* mice; n = 10/ Non-maternal-separated *App ^wt/wt^* mice, n = 13/ Non-maternal-separated *App^NL-G-F/wt^* mice, n = 8/ Maternal-separated *App ^wt/wt^* mice, n = 11/ Maternal-separated *App^NL-G-F/wt^* mice (Dunnett’s test). (C) Contextual memory and cued memory were measured using a fear-conditioning test. Behavioral layout of the fear conditioning test for the four experimental groups. Maternal-separated *App^NL-G-F/wt^* mice showed impairment of contextual memory. Contextual test: Main effect of group, F_(3,68)_ = 8.4507, p = 0.0001; main effect of time, F_(4,272)_ = 5.3835, p = 0.0003; interaction of group × time, F_(12,272)_ =1.0939, p = 0.3651; ** p < 0.01 Maternal-separated *App^NL-G-F/wt^* mice vs non-maternal-separated *App^wt/wt^* mice, *** p < 0.001 Maternal-separated *App^NL-G-F/wt^* mice vs non-maternal-separated *App^NL-G-F/wt^* mice; n = 19/ Non-maternal-separated *App ^wt/wt^* mice, n = 24/ Non-maternal-separated *App^NL-G-F/wt^* mice, n = 10/ Maternal-separated *App ^wt/wt^* mice, n = 22/ Maternal-separated *App^NL-G-F/wt^* mice (two-way repeated ANOVA followed by Holm’s test). (D) The formation of amyloid beta plaque detected by 82E1-biotin (cyan). The formation of amyloid beta plaque was facilitated by maternal separation in maternal-separated *App^NL-G-F/wt^* mice. *** p < 0.001 Maternal-separated *App^NL-G-F/wt^* mice vs other groups n = 4/each group (Tukey-Kramer test). Scale bar = 100 *μ*m.

### Maternal separation induced impairment of cognitive function and facilitated formation of amyloid beta plaque in the adult App^NL-G-F/wt^ mice

Figure 1B illustrates the effect of maternal separation on object location test which evaluate spatial memory (Denninger et al., 2018). Object location memory in the *App^wt/wt^* mice was not altered by maternal separation, as indicated by the fact that the discrimination index was not changed by maternal separation. By contrast, object location memory in the *App^NL-G-F/wt^* mice was impaired by maternal separation, as indicated by a discrimination index close to zero. The non-maternal-separated *App^NL-G-F/wt^* mice showed intact object location memory. These results indicate that the combination of APP mutation and maternal separation influence object location memory. Figure 1C indicates the effect of maternal separation on contextual and cued fear memory. Contextual fear memory in the *App^wt/wt^* mice was not altered by maternal separation, while that in the *App^NL-G-F/wt^* mice was impaired by maternal separation, as indicated by the lower freezing time in the maternal-separated *App^NL-G-F/wt^* mice in comparison with the non-maternal-separated *App^NL-G-F/wt^* and non-maternal separated *App^wt/wt^* mice. By contrast, tone-cued fear memory was not impaired in any of the four groups, indicating that contextual memory, which is related to prefrontal cortex–hippocampus signaling and is known to be necessary for the encoding of spatial cues, was impaired (Kitamura et al., 2017). Figure 1D indicates the effect of maternal separation on the formation of amyloid beta plaque. The formation of amyloid beta plaque in the *App^NL-G-F/wt^* mice was facilitated in mice that underwent maternal separation than in non-maternal-separated *App^NL-G-F/wt^* mice.

### Maternal separation induces vascular system impairment in adult mice

Phenotypes showing decreased pericyte coverage, a narrowing of capillary diameter, and impairment of the BBB are frequently observed in the postmortem brain of AD patients (Halliday et al., 2016). Figure 2 illustrates the effect of maternal separation on the vascular system. The maternal-separated *App^wt/wt^* and maternal-separated *App^NL-G-F/wt^* mice showed lower pericyte coverage of capillaries in the prefrontal cortex (PFC) than the non-maternal-separated controls in adult. Furthermore, maternal separation induced a narrowing of capillary diameter in both the *App^wt/wt^* mice and *App^NL-G-F/wt^* mice (Figure 2A). The more vessel-associated microglia were detected in the maternal-separated *App^NL-G-F/wt^* mice (Figure 2B). The result suggests strengthening of the interaction between microglia and vessels in the maternal-separated *App^NL-G-F/wt^* mice. The vessel-associated microglia migrate towards cerebral vessel and cause increased BBB permeability; Microglia, which can acquire a reactive phenotype that phagocytoses BBB components, initiate leakage of systemic substances into the parenchyma (Haruwaka et al., 2019). In our study, extra-vascular accumulation of fibrinogen deposits in the PFC was detected only in the maternal-separated *App^NL-G-F/wt^* mice (Figure 2C). We speculate that this accumulation was caused by leakage of fibrinogen due to impairment of the vascular system by more cessel-associated phagocytotic microglia only in the maternal-separated *App^NL-G-F/wt^* mice. Figure 2D shows the relationship between the sizes of the amyloid beta plaques and the distances from the amyloid beta plaques to vessels in the maternal-separated and non-maternal-separated *App^NL-G-F/wt^* mice that had mild amyloid beta plaque symptoms. The correlations between the sizes of amyloid beta plaques and the distances from the amyloid beta plaques to vessels were significant, which suggests the possibility that amyloid beta plaques were generated around the vessels and caused disappearance of the vessels after disruption. We also detected ameboid microglia surrounding amyloid beta plaques stained with tomato-lectin (Acarin et al., 1994).

**Figure 2.**
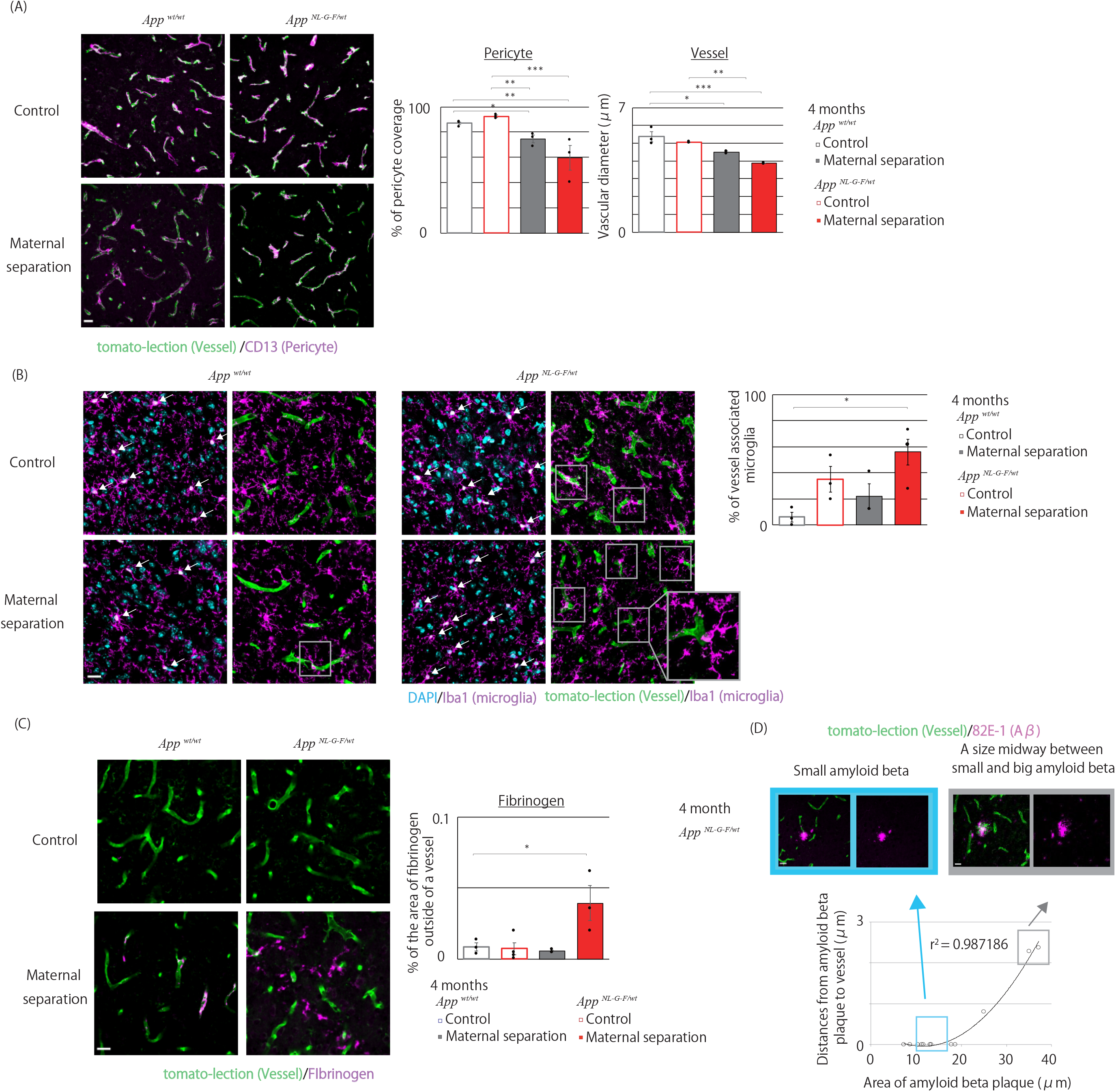
Maternal separation induced impairment to the vascular system. (A) The health of the vascular system was evaluated by tomato-lectin labelling of endothelial cells (green) and CD13 labelling of pericytes (magenta). Maternal separation induced lower pericyte coverage in *App^wt/wt^* mice and *App^NL-G-F/wt^* mice. * p < 0.05, Maternal-separated *App^wt/wt^* vs non-maternal-separated *App^wt/wt^* mice, ** p < 0.01 Maternal-separated *App^NL-G-F/wt^* vs non-maternal-separated *App^wt/wt^* mice, ** p < 0.01, Maternal-separated *App^wt/wt^* vs non-maternal-separated *App^NL-G-F/wt^* mice, *** p < 0.001 Maternal-separated *App^NL-G-F/wt^* vs non-maternal-separated *App^NL-G-F/wt^* mice. Furthermore, maternal separation also induced a narrowing of vascular diameter in both phenotypes. * p < 0.05 Maternal-separated *App^wt/wt^* mice vs non-maternal-separated *App^wt/wt^* mice, *** p < 0.001 Maternal-separated *App^NL-G-F/wt^* mice vs non-maternal-separated *App^wt/wt^* mice, ** p < 0.01 Maternal-separated *App^NL-G-F/wt^* mice vs non-maternal-separated *App^NL-G-F/wt^* mice; n = 3/each group (Tukey-Kramer test). Scale bar = 20 *μ* m. (B) Microglia (White arrow in Left image) were detected by microglial labelling with Iba1 (magenta) and DAPI (cyan). Vessel-associated microglia (Grey square in Right image) were detected by microglial labelling with Iba1 (magenta) and endothelial cell labelling with tomato-lectin (green). In maternal-separated *App^NL-G-F/wt^* mice, vessel associated microglia were increased. * p < 0.05 Maternal-separated *App^NL-G-F/wt^* mice vs non-maternal-separated *App^wt/wt^* mice; n = 3/ Non-maternal-separated *App ^wt/wt^* mice, n = 3/ Non-maternal-separated *App^NL-G-F/wt^* mice, n =3/ Maternal-separated *App ^wt/wt^* mice, n =4/ Maternal-separated *App^NL-G-F/wt^* mice (Dunnett’s test). (C) The integrity of the BBB was evaluated by tomato-lectin labelling of endothelial cells (green) and fibrinogen (magenta). Maternal separation caused accumulation of fibrinogen outside of vessels in maternal-separated *App^NL-G-F/wt^* mice. * p < 0.05 Maternal-separated *App^NL-G-F/wt^* mice vs non-maternal-separated *App^wt/wt^* mice; n = 3/each group (Dunnett test). Scale bar = 20 *μ*m. (D) The distance from the amyloid beta plaques to vessels and the area of amyloid beta plaque were determined using 82E1-biotin labelling of amyloid beta (magenta) and tomato-lectin labelling of endothelial cells (green). The area of amyloid beta plaque showed a significant association with the distance from the amyloid beta plaque to vessels in non-maternal-separated and maternal-separated *App^NL-G-F/wt^* mice. Small amyloid beta plaques are indicated with cyan squares, and amyloid beta plaques with a size midway between small and big amyloid beta plaques are indicated by grey squares. r^2=^ 0.987186 p < 0.001; n = 12 (polynomial regression). Scale bar = 20 *μ* m.

### Either maternal separation or APP mutation caused activation of and morphological change in microglia in adult mice

As the vascular system is affected by microglia, we examined the activity and morphology of microglia. Figure 3 shows the effects of maternal separation on microglia. Maternal separation caused activation of the microglia in the PFC of both *App^wt/wt^* mice and *App^NL-G-F/wt^* mice (Figure 3A). Although the numbers of microglia were not significantly different between any of the groups, except for the microglia surrounded by amyloid beta plaque, the proportions of soma and dendrites stained by Iba1 were lower in both *App^wt/wt^* mice and *App^NL-G-F/wt^* mice subjected to maternal separation (Figure 3A) than in the corresponding controls. Maternal separation may have induced morphological changes in both genotypes. More detailed analysis of the morphological changes revealed differences between non-maternal-separated *App^wt/wt^* mice and other groups (Figure 3B). Lower numbers of dendrite terminals and a shortening of the total branch length were detected in the maternal-separated *App^NL-G-F/wt^* mice than in the non-maternal-separated *App^wt/wt^* mice. On the other hand, the number of dendrite terminals was lower in the non-maternal-separated *App^NL-G-F/wt^* mice than in the non-maternal-separated *App^wt/wt^* mice (Figure 3B). The non-maternal-separated *App^NL-G-F/wt^* mice may have milder microglial symptoms than the maternal-separated animals. Furthermore, the circularity and solidity of microglia increased in only the maternal-separated *App^NL-G-F/wt^* mice. We speculated that maternal separation in combination with the APP mutation induced ameboid-like severe morphological change.

**Figure 3.**
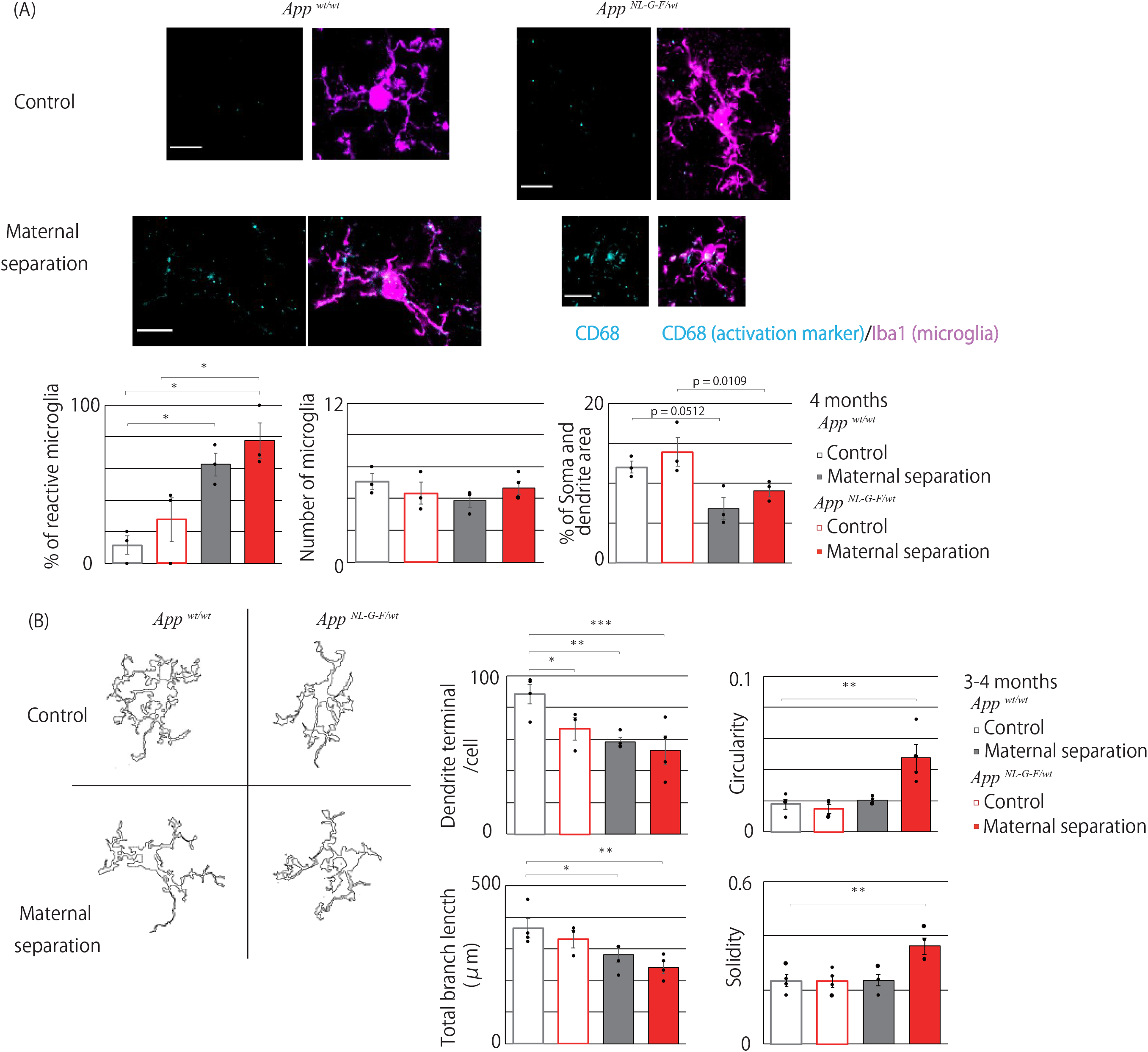
Maternal separation and genetic factors activated and changed the morphology of microglia. (A) Microglial morphology and activation were detected by labelling microglia with Iba1 (magenta) and labelling activation with CD68 (cyan). Maternal separation increased in activated microglia in *App^wt/wt^* and *App^NL-G-F/wt^* mice. * p < 0.05 Maternal-separated *App^wt/wt^* mice vs non-maternal-separated *App^wt/wt^* mice, * p < 0.05 Maternal-separated *App^NL-G-F/wt^* mice or non-maternal-separated *App^wt/wt^* mice. * p < 0.05 Maternal-separated *App^NL-G-F/wt^* mice or non-maternal-separated *App^NL-G-F/wt^* mice (Tukey-Kramer test). Furthermore, occupancy of soma and dendrites in maternal-separated *App^wt/wt^* and maternal-separated *App^NL-G-F/wt^* mice was lower. n = 3/ Non-maternal-separated *App ^wt/wt^* mice, n = 3/ Non-maternal-separated *App^NL-G-F/wt^* mice, n =3/ Maternal-separated *App ^wt/wt^* mice, n =4/ Maternal-separated *App^NL-G-F/wt^* mice (Tukey-Kramer test). Scale bar = 10 *μ*m. (B) Detailed morphological analysis was performed using the ImageJ plugin AnalyzeSkeleton. Images show examples of the outlines of microglia. Maternal separation and APP mutation decreased dendritic terminals in *App^wt/wt^* and *App^NL-G-F/wt^* mice. * p < 0.05 Non-maternal-separated *App^NL-G-F/wt^* mice vs non-maternal-separated *App^wt/wt^* mice, ** p < 0.01 Maternal-separated *App^wt/wt^* mice vs non-maternal-separated *App^wt/wt^* mice, ***p < 0.001 Maternal-separated *App^NL-G-F/wt^* mice vs non-maternal-separated *App^wt/wt^* mice (Dunnett test). Furthermore, the total branch length in maternal-separated *App^NL-G-F/wt^* mice and maternal-separated *App^wt/wt^* mice were shorter. * p < 0.05 Maternal-separated *App^wt/wt^* mice vs non-maternal-separated *App^wt/wt^* mice, ** p < 0.01 Maternal-separated *App^NL-G-F/wt^* mice vs non-maternal-separated *App^wt/wt^* mice. The circularity of microglia increased in only the maternal-separated *App^NL-G-F/wt^* mice. ** p < 0.01 Maternal-separated *App^NL-G-F/wt^* mice vs non-maternal-separated *App^wt/wt^* mice. The solidity of microglia increased in only the maternal-separated *App^NL-G-F/wt^* mice. ** p < 0.01 Maternal-separated *App^NL-G-F/wt^* mice vs non-maternal-separated *App^wt/wt^* mice; n =4/each group (Dunnett test).

### The maternal-separated App^wt/wt^ and App^NL-G-F/wt^ mice showed intact object location memory, absence of amyloid beta plaque, and normal vascular morphology and pericyte in adolescence

To detect initial changes in the maternal-separated and/or mutated mice, we examined the phenotypes at an earlier stage (P90), that is, in adolescence. Figure 4A shows the effect of maternal separation on object location memory in adolescence. In all groups, the discrimination index for object location memory indicated object location memory to be intact. Figure 4B shows that amyloid beta plaque was absent in all groups. Figure 4C shows no impairment of vascular morphology or pericytes in any of the groups. We detect no changes in cognitive function, plaque formation and vascular system in the maternal-separated and/or mutated mice at adolescence, which allow us to rule out that the maternal-separated and/or mutated mice are model mice for developmental disorders.

**Figure 4.**
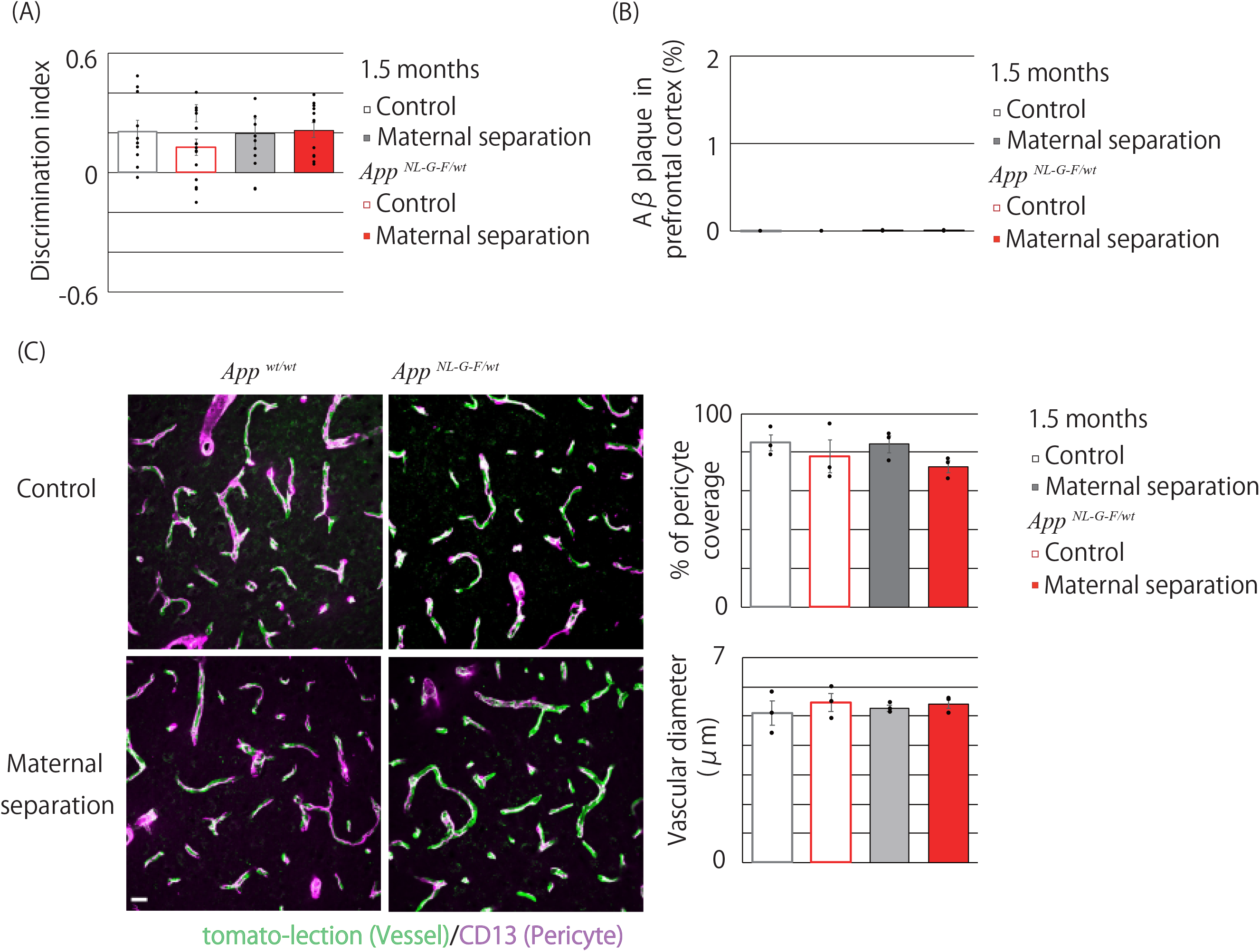
Alzheimer’s disease-like phenotypes did not begin during adolescence in maternal-separated App^NL-G-F/wt^ mice, and impairment of the vascular system did not begin during adolescence in all groups. (A) Spatial memory was measured using an object location test. Maternal-separated *App^NL-G-F/wt^* mice showed normal cognitive function. n = 9/ Non-maternal-separated *App ^wt/wt^* mice, n = 11/ Non-maternal-separated *App^NL-G-F/wt^* mice, n = 12/ Maternal-separated *App ^wt/wt^* mice, n = 11/ Maternal-separated *App^NL-G-F/wt^* mice (B) The formation of amyloid beta plaque detected by 82E1-biotin labelling of amyloid beta. No amyloid beta plaque was detected in maternal-separated *App^NL-G-F/wt^* mice. n = 3/each group (C) The health of the vascular system was evaluated by tomato-lectin labelling of endothelial cells (green) and CD13 labelling of pericytes (magenta). All groups showed normal pericyte coverage and a healthy vascular diameter. n = 3/each group. Scale bar = 20 *μ*m.

### Maternal separation and APP mutation cause functional and morphological change to microglia in adolescence

Figure 5 indicates the effect of maternal separation on microglia in adolescence. The numbers of activated microglia were not significantly different between each group (Figure 5A). A more detailed analysis of morphological differences revealed lower numbers of dendrite terminals in the maternal-separated *App^NL-G-F/wt^* mice than in the non-maternal-separated *App^wt/wt^* mice. In the *App^wt/wt^* mice, the maternally separated mice also showed lower numbers of dendrite terminals. Furthermore, lower numbers of dendrite terminals were detected in the non-maternal-separated *App^NL-G-F/wt^* mice than in the non-maternal-separated *App^wt/wt^* mice (Figure 5B). On the other hand, the numbers of vessel-associated microglia did not differ between the groups (Figure 5C). To evaluate the pro-inflammatory properties of microglia, we measured the level of TNF-R1 protein in the PFC. Maternal-separated *App^wt/wt^* and *App^NL-G-F/wt^* mice showed higher levels of TNF-R1 protein than the non-maternal-separated mice (Figure 5D). We detected changes associated with both maternal separation and heterozygotic mutation in adolescence, indicating the possibility that a change in microglia caused by both maternal separation and APP mutation induces an AD-like phenotype. To assess the effect of maternal separation on the hypothalamus-pituitary-adrenal axis in adolescence, we measured serum corticosterone levels induced by a novel cage stress in maternal-separated and non-separated mice. Maternal-separated *App^NL-G-F/wt^* and *App^wt/wt^* mice subjected to a novel cage stress in adolescence mice showed more corticosterone release than non-maternal-separated *App^NL-G-F/wt^* and *App^wt/wt^* mice (Figure 5E).

**Figure 5.**
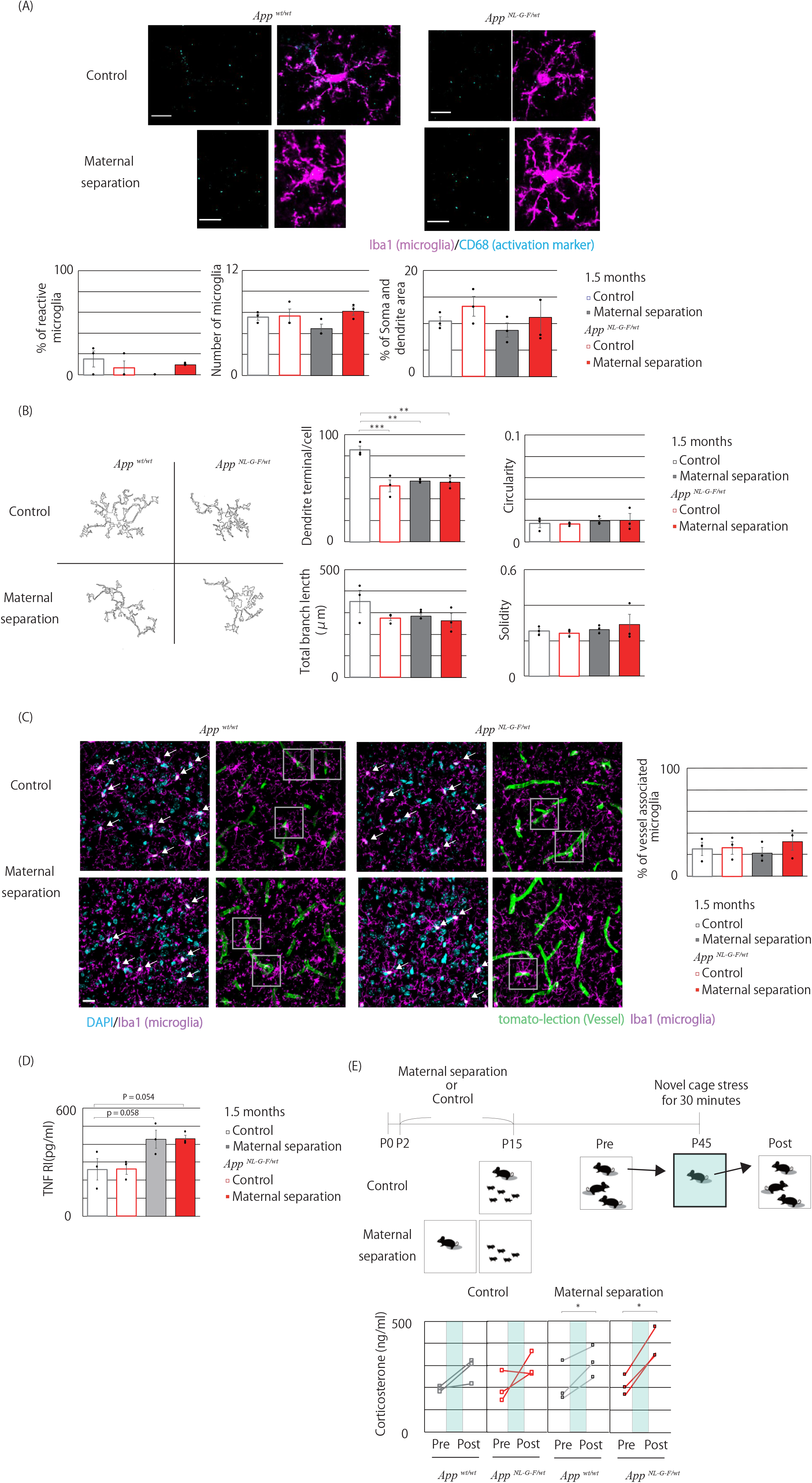
Maternal separation and genetic factors induced morphological changes in microglia and maternal separation in combination with genetic factors led to abnormality in the stress response during adolescence. (A) Microglial morphology and activation were detected by labelling with Iba1 (magenta) and CD68 (cyan), respectively. Increased activation of microglia was not observed in maternal-separated *App^wt/wt^* and maternal-separated *App^NL-G-F/wt^* mice. n = 3/each group. Scale bar = 10 *μ* m. (B) Detailed morphological analysis was performed using the ImageJ. Maternal separation and APP mutation decreased dendritic terminals in *App^wt/wt^* and *App^NL-G-F/wt^* mice. *** p < 0.001 Non-maternal-separated *App^NL-G-F/wt^* mice vs non-maternal-separated *App^wt/wt^* mice, ** p < 0.01 Maternal-separated *App^wt/wt^* mice vs non-maternal-separated *App^wt/wt^* mice, ** p < 0.01 Maternal-separated *App^NL-G-F/wt^* mice vs non-maternal-separated *App^wt/wt^* mice (Dunnett test). (C) Microglia (white arrow in left image) were detected by microglial labelling with Iba1 (magenta) and DAPI (cyan). Vessel-associated microglia (Grey square in right image) were detected by microglial labelling with Iba1 (magenta) and endothelial cell labelling with tomato-lectin (green). The numbers of vessel-associated microglia were similar across the four experimental groups. n = 3/each group. Scale bar = 20 *μ*m. (D) The protein level of TNF-R1 was higher in maternal-separated *App^wt/wt^* and maternal-separated *App^NL-G-F/wt^* mice. p = 0.058 Maternal-separated *App^wt/wt^* mice vs non-maternal-separated *App^wt/wt^* mice, p = 0.054 Maternal-separated *App^NL-G-F/wt^* mice vs non-maternal-separated *App^wt/wt^* mice n = 3/each group (Dunnett test). (E) The response to novel cage stress was measured to assess an effect of maternal separation on the HPA axis. Not only *App^NL-G-F/wt^* mice but also *App^wt/wt^* mice showed that increased stress caused by maternal separation induced corticosterone release. Maternal-separated *App^NL-G-F/wt^* mice: F (1,2) = 75.414, p = 0.0130; Maternal-separated *App^wt/wt^* mice: F _(1,2)_ = 22.035, p = 0.0425. * p < 0.05 vs corticosterone before novel cage stress; n = 3/each group (Paired t-test).

## Discussion

In this study, we investigated whether maternal separation facilitates the development of AD, with the aim of understanding the causal association between early-life stress and the development and progression of AD. We also investigated for changes induced by maternal separation, particularly in capillary vessels. We established that maternal separation facilitates the impairment of spatial cognitive function and the formation of amyloid beta plaque in *App^NL-G-F/wt^* mice, with disruption of micro-capillaries, and we verified that early-life stress constitutes a risk factor for AD. Furthermore, we found that morphological and functional changes to microglia are early symptoms in our experimental model, and suggest the possibility that impairment of the cerebral vascular system through microglia caused by combination of maternal separation and APP mutation induces dysfunction in the BBB, thereby facilitating the clinical condition of AD.

Research using the Childhood Trauma Questionnaire to assess childhood stress has suggested that early-life stress may be associated with increased risk of AD and dementia (Radford et al., 2017). However, because most previous reports, including the above research, were based on retrospective cohort studies, it remains unclear whether early-life stress affects the onset and progression of AD. Although it has been reported that maternal separation facilitates the formation of amyloid beta plaque using an AD mouse model different from the one we used (Hui et al., 2017), the pathway by which maternal separation induces increased amyloid beta plaque remains unclear. We found that maternal separation facilitates the impairment of spatial cognitive function and the formation of amyloid beta plaque through impairment of the cerebral vascular system in *App^NL-G-F/wt^* mice. We propose that these maternally separated *App^NL-G-F/wt^* mice can be used as a stress-induced AD mouse model. As it is known that progression of AD in humans is exacerbated by stress, such as a change in abode and hospital admission in old age, our AD mouse model may contribute to understanding the mechanism of stress-induced AD and how it can be prevented.

We found that maternal separation mildly impaired the cerebral capillary vascular system. However, dysfunction of the BBB, which is a more serious symptom, was detected in the maternal-separated *App^NL-G-F/wt^* mice and was related to both maternal separation and genetic factors. Several mechanisms are involved in clear amyloid beta clearance (Chen et al., 2017). One mechanism involves transcytosis through cells of the BBB. Another mechanism involves phagocytosis by microglia. The reduction in amyloid beta clearance due to impairment of the vascular system and impaired microglial phagocytoses of amyloid beta may induce inflammation and increase fibrin generation due to coagulation factor XII (FXII) activation by amyloid beta. The interaction of amyloid beta with fibrin (or fibrinogen) can lead to increased fibrin deposition in cerebral blood vessels, and these accumulated fibrin deposits may induce microinfarcts, which result in hemorrhage, inflammation, and BBB disruption, all of which are commonly observed in AD (Ahn et al., 2017). Our result that disappearance of vessel around larger amyloid plaque (Fig. 2D) may be caused by the interaction of amyloid beta with fibrin.

We report that maternal separation induces impairments to the cerebral vascular system, with impairments to pericytes and narrowing of vessels in adulthood. Both of these phenotypes are detected in the postmortem brains of human patients with AD (Halliday et al., 2016). Therefore, these findings indicate a relationship between early-life stress and AD pathology. It has been demonstrated that both antenatal and postnatal stress delay the development of the BBB during weaning (Gomez-Gonzalez and Escobar, 2009). We detected no impairment in vascular morphology or in pericytes in adolescence. Our findings that early-life stress induces impairments in adult mice is in accord with the Development Origins of Health and Disease (DOHaD) hypothesis, which purports that exposure to certain environmental conditions during critical periods of development and growth may have significant adverse effects on short- and long-term health. As the ultrastructural breakdown of capillaries and a decrease in blood flow are detected in AD patients and AD mouse models (de la Torre, 2004), it is likely that impairment of the cerebral vascular system develops before the onset of AD pathogeny in *App^NL-G-F/wt^* mice.

We detected morphological changes in microglia and changes in their pro-inflammatory properties (which are mediated by the activation of TNF-R1) at an early stage, while we did not detect amyloid plaque formation, changes to the vascular system, or changes in behavior. Microglia have been reported to show morphological changes in response to the clinical condition of AD (Yamada and Jinno, 2013). An increase in corticosterone in response to stress has been reported to be related to a change in microglia (Rincon-Cortes and Sullivan, 2014). In our study, maternal-separated *App^NL-G-F/wt^* and maternal-separated *App^wt/wt^* mice showed increased corticosterone release in response to a novel cage stress at adolescence, which suggests that maternal separation affects the stress response. In fact, the production of corticosterone in response to stress facilitates the release of monocytes into the circulation from bone marrow, which in turn facilitates monocyte adherence to vessels (Niraula et al., 2018). The release of monocytes upregulates IL-1β and stimulates endothelial cells (McKim et al., 2018). Haruwaka et al. hypothesized that endothelial cells may respond to systemic inflammation and release molecules to trigger microglia migration and phenotype changes. They also reported that chemokine (C-C motif) ligand 5 (CCL5), which is secreted by endothelial cells, attracts microglia to vessels. Recently, plasma from aged mice was shown to affect the activity of microglia, referred to as neuroinflammation, and increase VCAM1 expression on endothelial cell, a major cell type of the BBB, in young mice (Yousef et al., 2019). The authors suggested that circulating cytokines and chemokines with detrimental effects on the brain give rise to these changes in microglia and endothelial cells. Considering that increased age is crucial for the development of AD, increases in inflammatory signals in the plasma may induce AD. On the other hand, the non-maternal-separated *App^NL-G-F/wt^* mice also showed morphological changes in adolescence. Our data support the hypothesis that impairment of the vascular system by amyloid beta oligomers induces activation of microglia (Nortley et al., 2019). As microglia and the vascular system are deeply linked, the possibility that an undetectable alteration of the vascular system led to a change in the microglia, which then led to the development of AD via an interaction between microglia and the vascular system, cannot be ruled out. The changes in microglia induced by maternal separation and those induced by APP mutation may occur via different mechanisms. Microglia activation induced by maternal separation in combination with *App^NL-G-F/wt^* mice may impairs the vascular system, leading to AD progression.

In the present study, we established that maternal separation facilitates the development of AD. Furthermore, we identified changes in microglia induced by maternal separation and/or genetic factors as an early phenotype, which probably preceded the impairment of the vascular system in the stress-induced AD mouse model. These results indicate that microglia could have potential use as biomarkers and as targets for new therapies against AD.

## Author contributions

T.T. and H.O. designed the research. T.T. performed and analyzed all experiments. T.S. and T.S. generated and provided App mutant mice (*App^NL-G-F/NL-G-F^*). S.H., M.H., H.S., and M.H. helped with this study and made important suggestions. T.T. generated all the figures and tables and wrote the manuscript. H.O. edited the manuscript and supervised the study.

## Acknowledgments

The authors thank Dr. Makoto Hashimoto for advice. We would like to thank Bioedit Ltd (https://www.bioedit.com) for English language editing. This work was partially supported by JSPS KAKENHI Grant Number 18H02537 for Haruo Okado, and by JSPS KAKENHI Grant Number 17K16408 and Research Grant for Public Health Science for Tomoko Tanaka.

## References

Acarin, L., Vela, J.M., González, B., Castellano, B., 1994. Demonstration of poly-N-acetyl lactosamine residues in ameboid and ramified microglial cells in rat brain by tomato lectin binding. J. Histochem. Cytochem. 42, 1033–1041.

Ahn, H.J., Chen, Z.L., Zamolodchikov, D., Norris, E.H., Strickland, S., 2017. Interactions of β-amyloid peptide with fibrinogen and coagulation factor XII may contribute to Alzheimer’s disease. Curr. Opin. Hematol. 24, 427–431.

Chen, G.F., Xu, T.H., Yan, Y., Zhou, Y.R., Jiang, Y., Melcher, K., Xu, H.E., 2017. Amyloid beta: structure, biology and structure-based therapeutic development. Acta Pharmacol. Sin. 38, 1205–1235.

Christensen, K., Johnson, T.E., Vaupel, J.W., 2006. The quest for genetic determinants of human longevity: challenges and insights. Nature reviews. Genetics 7, 436–448.

de la Torre, J.C., 2004. Is Alzheimer’s disease a neurodegenerative or a vascular disorder? Data, dogma, and dialectics. Lancet Neurol. 3, 184–190.

Denninger, J.K., Smith, B.M., Kirby, E.D., 2018. Novel Object Recognition and Object Location Behavioral Testing in Mice on a Budget. J Vis Exp.

Fernández-Arjona, M.D.M., Grondona, J.M., Granados-Durán, P., Fernández-Llebrez, P., López-Ávalos, M.D., 2017. Microglia Morphological Categorization in a Rat Model of Neuroinflammation by Hierarchical Cluster and Principal Components Analysis. Front. Cell. Neurosci. 11, 235.

Gomez-Gonzalez, B., Escobar, A., 2009. Altered functional development of the blood-brain barrier after early life stress in the rat. Brain Res. Bull. 79, 376–387.

Halliday, M.R., Rege, S.V., Ma, Q., Zhao, Z., Miller, C.A., Winkler, E.A., Zlokovic, B.V., 2016. Accelerated pericyte degeneration and blood-brain barrier breakdown in apolipoprotein E4 carriers with Alzheimer’s disease. J. Cereb. Blood Flow Metab. 36, 216–227.

Haruwaka, K., Ikegami, A., Tachibana, Y., Ohno, N., Konishi, H., Hashimoto, A., Matsumoto, M., Kato, D., Ono, R., Kiyama, H., Moorhouse, A.J., Nabekura, J., Wake, H., 2019. Dual microglia effects on blood brain barrier permeability induced by systemic inflammation. Nat Commun 10, 5816.

Hui, J., Feng, G., Zheng, C., Jin, H., Jia, N., 2017. Maternal separation exacerbates Alzheimer’s disease-like behavioral and pathological changes in adult APPswe/PS1dE9 mice. Behav. Brain Res. 318, 18–23.

Iadecola, C., 2017. The Neurovascular Unit Coming of Age: A Journey through Neurovascular Coupling in Health and Disease. Neuron 96, 17–42.

Kitamura, T., Ogawa, S.K., Roy, D.S., Okuyama, T., Morrissey, M.D., Smith, L.M., Redondo, R.L., Tonegawa, S., 2017. Engrams and circuits crucial for systems consolidation of a memory. Science 356, 73–78.

Mawuenyega, K.G., Sigurdson, W., Ovod, V., Munsell, L., Kasten, T., Morris, J.C., Yarasheski, K.E., Bateman, R.J., 2010. Decreased clearance of CNS beta-amyloid in Alzheimer’s disease. Science 330, 1774.

McKim, D.B., Weber, M.D., Niraula, A., Sawicki, C.M., Liu, X., Jarrett, B.L., Ramirez-Chan, K., Wang, Y., Roeth, R.M., Sucaldito, A.D., Sobol, C.G., Quan, N., Sheridan, J.F., Godbout, J.P., 2018. Microglial recruitment of IL-1beta-producing monocytes to brain endothelium causes stress-induced anxiety. Mol.Psychiatry 23, 1421–1431.

Montagne, A., Nation, D.A., Pa, J., Sweeney, M.D., Toga, A.W., Zlokovic, B.V., 2016. Brain imaging of neurovascular dysfunction in Alzheimer’s disease. Acta Neuropathol. 131, 687–707.

Niraula, A., Wang, Y., Godbout, J.P., Sheridan, J.F., 2018. Corticosterone Production during Repeated Social Defeat Causes Monocyte Mobilization from the Bone Marrow, Glucocorticoid Resistance, and Neurovascular Adhesion Molecule Expression. J. Neurosci. 38, 2328–2340.

Nortley, R., Korte, N., Izquierdo, P., Hirunpattarasilp, C., Mishra, A., Jaunmuktane, Z., Kyrargyri, V., Pfeiffer, T., Khennouf, L., Madry, C., Gong, H., Richard-Loendt, A., Huang, W., Saito, T., Saido, T.C., Brandner, S., Sethi, H., Attwell, D., 2019. Amyloid beta oligomers constrict human capillaries in Alzheimer’s disease via signaling to pericytes. Science 365.

Norton, M.C., Smith, K.R., Ostbye, T., Tschanz, J.T., Schwartz, S., Corcoran, C., Breitner, J.C., Steffens, D.C., Skoog, I., Rabins, P.V., Welsh-Bohmer, K.A., 2011. Early parental death and remarriage of widowed parents as risk factors for Alzheimer disease: the Cache County study. Am. J. Geriatr. Psychiatry 19, 814–824.

Radford, K., Delbaere, K., Draper, B., Mack, H.A., Daylight, G., Cumming, R., Chalkley, S., Minogue, C., Broe, G.A., 2017. Childhood Stress and Adversity is Associated with Late-Life Dementia in Aboriginal Australians. Am. J. Geriatr. Psychiatry 25, 1097–1106.

Rattray, I., Scullion, G.A., Soulby, A., Kendall, D.A., Pardon, M.C., 2009. The occurrence of a deficit in contextual fear extinction in adult amyloid-over-expressing TASTPM mice is independent of the strength of conditioning but can be prevented by mild novel cage stress. Behav. Brain Res. 200, 83–90.

Rincon-Cortes, M., Sullivan, R.M., 2014. Early life trauma and attachment: immediate and enduring effects on neurobehavioral and stress axis development. Front. Endocrinol. (Lausanne) 5, 33.

Saito, T., Matsuba, Y., Mihira, N., Takano, J., Nilsson, P., Itohara, S., Iwata, N., Saido, T.C., 2014. Single App knock-in mouse models of Alzheimer’s disease. Nat. Neurosci. 17, 661–663.

Seifan, A., Schelke, M., Obeng-Aduasare, Y., Isaacson, R., 2015. Early Life Epidemiology of Alzheimer’s Disease - A Critical Review. Neuroepidemiology 45, 237–254.

Sweeney, M.D., Ayyadurai, S., Zlokovic, B.V., 2016. Pericytes of the neurovascular unit: key functions and signaling pathways. Nat. Neurosci. 19, 771–783.

Wen, Y., Yang, S., Liu, R., Simpkins, J.W., 2004. Transient cerebral ischemia induces site-specific hyperphosphorylation of tau protein. Brain Res. 1022, 30–38.

Xu, H., Rajsombath, M.M., Weikop, P., Selkoe, D.J., 2018. Enriched environment enhances β-adrenergic signaling to prevent microglia inflammation by amyloid-β. EMBO Mol. Med. 10.

Yamada, J., Jinno, S., 2013. Novel objective classification of reactive microglia following hypoglossal axotomy using hierarchical cluster analysis. J. Comp. Neurol. 521, 1184–1201.

Yousef, H., Czupalla, C.J., Lee, D., Chen, M.B., Burke, A.N., Zera, K.A., Zandstra, J., Berber, E., Lehallier, B., Mathur, V., Nair, R.V., Bonanno, L.N., Yang, A.C., Peterson, T., Hadeiba, H., Merkel, T., Korbelin, J., Schwaninger, M., Buckwalter, M.S., Quake, S.R., Butcher, E.C., Wyss-Coray, T., 2019. Aged blood impairs hippocampal neural precursor activity and activates microglia via brain endothelial cell VCAM1. Nat. Med. 25, 988–1000.

Zlokovic, B.V., 2011. Neurovascular pathways to neurodegeneration in Alzheimer’s disease and other disorders. Nat. Rev. Neurosci. 12, 723–738.

